# The Opposing Effect of Hedonic and Eudaimonic Happiness on Gene Expression is Correlated Noise

**DOI:** 10.1101/044917

**Authors:** Jeffrey A. Walker

## Abstract

**Background:** This paper presents a re-analysis of the gene set data from Fredrickson et al. 2013 and Fredrickson et al. 2015 which purportedly showed opposing effects of hedonic and eudaimonic happiness on the expression levels of a set of genes that have been correlated with social adversity. Fredrickson et al. 2015 used a linear model of fixed effects with correlated error (using GLS) to estimate the partial regression coefficients.

**Methods:** The standardized effects of hedonic and eudaimonic happiness on CTRA gene set expression estimated by GLS was compared to estimates using multivariate (OLS) linear models and generalized estimating equation (GEE) models. The OLS estimates were tested using a bootstrap *t*-test, O’Brien’s OLS test, a permutation *t* test, and the rotation *z*-test. The GEE estimates were tested using a Wald test with robust standard errors. The performance (type I, type II, and type M error) of all tests was investigated using a Monte Carlo simulation of data modeled after the 2015 dataset.

**Results:** Standardized OLS effects (mean partial regression coefficients) of Hedonia and Eudaimonia on gene expression levels are very small in both the 2013 and 2015 data, as well as the combined data.The *p*-values from all tests fail to reject any of the null models. The GEE estimates and tests are nearly identical to the OLS estimates and tests. By contrast, the GLS estimates are inconsistent between data sets, but in each dataset, at least one coefficient is large and highly statistically significant. The Monte Carlo simulation of error rates shows inflated type I error from the GLS test on data with a similar correlation structure to that in the 2015 dataset, and this error rate increases as the number of outcomes increases relative to the number of subjects. Bootstrap and permutation GLS distributions suggest that the GLS model not only results in downward biased standard errors but also inflated coefficients. Both distributions also show the expected, strong, negative correlation between the coefficients for *Hedonia* and *Eudaimonia*.

**Discussion:** The results fail to support opposing effects, or any detectable effect, of hedonic and eudaimonic well being on the pattern of gene expression. The apparently replicated pattern of hedonic and eudaimonic effects on gene expression is most parsimoniously explained as "correlated noise" due to the geometry of multiple regression. A linear mixed model for estimating fixed effects in designs with many repeated measures or outcomes should be used cautiously because of the potentially inflated type 1 and type M error.

## Introduction

In a highly visible gene set analysis, Fredrickson et al. 2013 [8] claimed that a measure of eudaimonic happiness was associated with a decreased “conserved transcriptional response to adversity” (CTRA) while a measure of hedonic happiness was associated with increased CTRA. This transcriptional response includes the up-regulation of pro-inflammatory signals and the down-regulation of antiviral and antibody synthesis signals. Fredrickson et al. 2015 [7] followed up with a replicate study, found a large negative effect of *Eudaimonia* on CTRA expression, and emphasized that the opposing effects of hedonic and eudaimonic scores on mean CTRA gene expression replicated that of Fredrickson et al. 2013 [8].

Both the 2013 and 2015 analyses contain numerous statistical issues that question the conclusions. The initial (2013) data were analyzed using an *ad hoc* one sample *t*-test of the mean response. The *t*-test was implemented in a way that ignored the assumption of independent errors, resulting in inflated type 1 error, which was discovered by [3] via simulation. The replicate (2015) data were analyzed using a linear model with fixed-effects and correlated error (a type of linear mixed model although no random effect is explicitly modeled). In this replicate analysis, the authors failed to acknowledge the well known downward bias of standard errors when using this model with limited data [17].

Here, I present the results of a re-analysis of the Fredrickson et al. 2013 [8] and Fredrickson et al. 2015 [7] data. Not being a social psychologist, I limit my analysis to addressing the question “what is the evidence for effects of hedonic and eudaimonic happiness scores on CTRA gene expression” and give here only the necessary background to understand my analysis. The general question addressed by Fredrickson et al. 2013 [8] and Fredrickson et al. 2015 [7], that is, is there a mean response different from zero for a set of multiple outcomes, has a long and rich history in applied statistics [19], including in association studies of gene sets [12, 1]. [5] is an especially clear exposition of the different null hypotheses that one might test. Wu et al. [25] clearly outline some of these hypotheses in the context of gene set associations. Neither Fredrickson et al. 2013 [8] nor Fredrickson et al. 2015 [7] use any of these seminal papers to guide their analyses.

The datasets were re-analyzed using multivariate linear models (OLS) and generalized estimating equation models (GEE). The results from all analyses for each dataset are consistent in that they all suggest very small effects of *Hedonia* and *Eudaimonia* on CTRA gene expression and *p*-values that fail to provide evidence against the nulls. I also show that the high positive correlation between the *Hedonia* and *Eudaimonia* results in negatively correlated partial regression coefficients that may be misinterpreted as a “replicable” pattern instead of correlated noise reflecting the geometry of multiple regression.

A Monte-Carlo simulation experiment to model type I and II error rates shows excellent performance for the OLS and GEE models used in this study but inflated type I error for the linear mixed model (GLS) method used in Fredrickson et al. 2015 [7]. In addition, the Monte-Carlo simulation shows that the linear mixed model results in inflated coefficient estimates, or type M (magnitude) error [10], when the number of response variables increases relative to the number of subjects. Both type 1 and type M error decrease toward nominal levels with increased number of subjects. These Monte-Carlo results suggest caution when using linear mixed models to estimate fixed effects in designs with many repeated measures or multiple outcomes and also that Monte-Carlo simulations can be useful for testing model performance on simulated data modeled after empirical data.

### Background

The CTRA gene set includes 19 pro-inflammatory, 31 anti-viral, and 3 antibody-stimulating genes. The Fredrickson et al. 2013 [8] data included all 53 genes but the Fredrickson et al. 2015 [7] data is missing IL-6 from the pro-inflammatory subset.

Fredrickson et al. 2013 [8] used 53 univariate multiple regressions to estimate the effects (the regression coefficient) of each happiness (hedonic and eudaimonic) score on log2(normalized gene expression) for each gene. The regression model included both happiness scores, seven covariates to adjust for demographic and general health confounding (sex, age, ethnicity, BMI, a measure of alcohol consumption, a measure of smoking, and a measure of recent illness), and eight expression levels of T-lymphocyte markers to adjust for immune status confounding. Hedonic and eudaimonic scores were transformed to z-scores prior to the analysis. The 53 multiple regressions (one for each gene) yielded 53 coefficients for hedonic score and 53 coefficients for eudaimonic score. The coefficients of the 31 anti-viral and 3 antibody genes were multiplied by −1 to make the direction of the effect consistent with the CTRA response. Fredrickson et al. 2013 [8] used a simple one-sample *t*-test of the 53 coefficients to test for a mean effect of hedonic or eudaimonic score on CTRA expression. A mean coefficient greater than zero reflects a positive CTRA response (increased pro-inflammatory and decreased anti-viral and antibody-stimulating genes).

Fredrickson et al. 2013 [8] used a bootstrap to re-sample the coefficients in order to generate a standard error (the denominator of their *t*-value and then tested the statistic using *m* − 1 degrees of freedom, where *m* is the number of outcomes (gene expression levels). There are two fundamental problems with this *t*-test. First, the coefficients are not independent of each other because of the correlated expression levels among genes and as a consequence the standard error in the denominator will be too small, which should result in an inflated Type I error rate. Second, their degrees of freedom does not account for the number of subjects in the study. At the extreme, if only a single gene expression level is measured, Fredrickson’s *t* cannot even be computed. This second error should result in loss of power. The combined effect on Type I and Type II error will depend on the magnitude of the correlations among the expression levels. Through simulation, however, [3] discovered an inflated Type-I error in their exploration of the data using the Fredrickson et al. 2013 [8] *t*-test. [19] developed an appropriate *t*-test for the effects of an an independent variable on multiple outcomes (see below).

Fredrickson et al. 2015 [7] replicated the 2013 study but treated the 52 gene expression levels as “repeated” measures (or multiple outcomes) of a single expression response and used a linear model with fixed-effects and correlated error to estimate the regression coefficients of expression on hedonic and eudaimonic score. Specifically, Fredrickson et al. 2015 [7] used generalized least squares (GLS) with a heterogenous compound symmetry error matrix to estimate the marginal (population-averaged) fixed effects. Compound symmetry assumes equal correlation (conditional on the set of predictors) among all expression levels. This is not likely to approximate the true error structure for a set of expression levels for different genes, as these expression levels will share different sets of underlying regulatory factors. Fredrickson et al. 2015 [7] re-ran the analysis using an unstructured error matrix, with results contradicting the compound symmetry results, but chose to report this in the supplement and not the main text. While a GLS estimate of marginal effects is consistent if the error correlation is misspecified, the variance of the estimates is biased downwards, resulting in inflated Type I error [15, 13]. The amount of bias depends on the true and specified correlation structure, as well as effective sample size (a function of the number of subjects, the number of outcomes, and the correlations among the outcomes), but can be large even with large samples [13]. [17] further note that not only are the standard errors from the GLS model biased downwards but the coefficient estimates are biased upwards and that these behaviors increase with the complexity of the correlated error structure. When only the marginal effects are of interest (as here), population-averaged effects are typically estimated using Generalized Estimating Equations (GEE) instead of GLS and standard errors robust to model misspecification are computed using the sandwich estimator [16, 26].

## Methods

Data were downloaded as.txt Series Matrix Files from http://www.ncbi.nlm.nih.gov/geo/ using accession numbers GSE45330 (the 2013 dataset, hereafter FRED13) and GSE55762 (the 2015 dataset, hereafter FRED15). The CTRA (response) expression data were log2 transformed. The T-lymphocyte expression data that formed part of the set of covariates were log2 transformed in the downloaded data. The downloaded hedonic and eudaimonic scores in FRED13 had means and variances close but not equal to that expected of z-scores, which suggests that the public data slightly differs from that analyzed by Fredrickson et al. 2013 [8]; these were re-standardized to z-scores Three rows of FRED13 had missing covariate data (two rows were completely missing) and were excluded; the number of rows (cases) in the cleaned matrix was 76. The downloaded hedonic and eudaimonic scores in FRED15 were the raw values and were transformed to z-scores. There was no missing data in FRED15 and the number of cases was 122.

Prior to all analyses, *Hedonia* or *Eudaimonia* scores and the expression levels of all genes were standardized to mean zero and unit variance. Additionally, the 31 anti-viral and 3 antibody genes were multiplied by −1 to make the direction of the effect consistent with the CTRA response [8, 7].

Several judgements were necessary for the combined (FRED13+15) data. The covariate *illness* appears to represent a sum over 13 symptoms for the FRED13 data but an average for the FRED15 data. I only recover the Fredrickson et al. 2015 [7] results with the raw, downloaded data and not the FRED13 data averaged. For my analyses *illness* is left as is (as downloaded) to be consistent with the previously published results. For the standardized coefficients in the combined data, the raw values in each dataset should be combined first and then the entire column should be centered and scaled. But because the *Hedonia* and *Eudaimonia* variables are pre-standardized in the separate public datasets, the expression variables were standardized before combining but not after, in order to maintain the correct relationship between independent and dependent variables within each study of the combined data. This procedure also is consistent with previously published results. Finally, the multivariate methods that I use cannot handle missing data and, consequently, in order for me to compare analyses among inference methods, the response *IL*6 was dropped from the combined data since this was missing from FRED2015. This affected only the analysis of the combined data and had trivial effect on the coefficient estimates.

### Null hypothesis tests

If *β_j_* is the the effect (partial regression coefficient) of *Hedonia* or *Eudaimonia* on the expression level of the *j*th gene, the overall effect of *Hedonia* or *Eudaimonia* on expression of the CTRA gene set is simply the averaged coefficient over all genes, 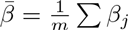 where *m* is the number of genes. The three focal null hypotheses that are tested here, which were also the focus of Fredrickson et al. 2013 [8] and Fredrickson et al. 2015 [7] are 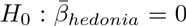,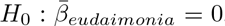, and 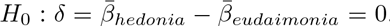. A11 three hypotheses are directional, that is, the mean effect differs from zero. This differs from the general multivariate test that at least one of the coefficients differs from zero, but the mean response may be zero. While the hypotheses are directional, the tests are two-tailed, that is, the mean response may be up or down regulation of the CTRA gene set.

### OLS inferential tests

The effects of *Hedonia* and *Eudaimonia* on the mean of the *m* gene expression levels are estimated with the multivariate linear model

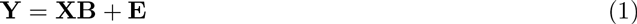
where **Y** is the *n × m* matrix of gene expression levels for the *n* subjects, **X** is the model matrix of dummy variables and covariates, **E** is the matrix of residual error, and **B** is the *p × m* matrix of partial regression coefficients. For the combined data, the model matrix includes a dummy variable indicating dataset (2013 or 2015). The coefficients of the *j*th column of **B** are precisely equal to univariate multiple regression of the *j*th gene on **X** (and why the model is sometimes called a multivariate multiple regression). In **R**, estimating the m effects of *Hedonia* and *Eudaimonia* is much faster using this multivariate model than looping through *m* univariate multiple regressions. The mean of the *m* coefficients is the OLS estimate of the effect of *Hedonia* or *Eudaimonia* on overall CTRA expression level. Because the happiness scores for *Hedonia* and *Eudaimonia* and the *m* expression levels were mean-centered and variance-standardized, the reported OLS estimates are mean (averaged over the *m* genes) standard partial regression coefficients.

### Bootstrap *t*-test

Fredrickson et al. 2013 [8] used a bootstrap resampling method to compute a *p*-value. In their bootstrap, the 53 partial regression coefficients were re-sampled with replacement 200 times. Each iteration, a mean regression coefficient was computed. The standard deviation of the 200 means was used as the estimate of the standard error to compute a *t*-statistic and associated *p*-value. Resampling the regression coefficients fails to address the lack of independence among the coefficients. To estimate the sampling error that accounts for correlated error among the regression coefficients, the entire estimation procedure needs to be included within the bootstrap by resampling the data and re-estimating the coefficients. In each iteration of this procedural bootstrap, entire rows of the data were re-sampled with replacement, the *m* coefficients were estimated by equation 1, and the re-sampled mean coefficients (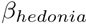 and 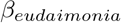) were saved each iteration. 1999 bootstrap iterations were run. Standard errors of the coefficients were computed as the standard deviation of the 2000 saved mean coefficients. Additionally, a *t* test in the spirit of Fredrickson et al. 2013 [8] was computed using the observed mean coefficient divided by its standard error as the *t*-value. Like that from Fredrickson et al. 2013 [8], this *t* fails to account for the multiple outcomes within subject design addressed by the other tests described below.

### O’Brien’s OLS *t*-test

O’Brien’s OLS test [19, 18, 6] was developed explicitly for testing the directional hypothesis that the mean effect of multiple outcomes differs from zero, which is precisely the question pursued in both Fredrickson papers. Given *m* standardized regression coefficients and associated *t*-values, O’Brien’s test statistic is

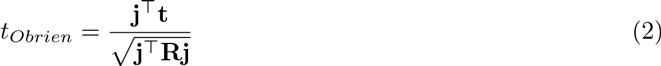

where **j** is a *m* vector of 1s, *t* contains the *t*-values associated with each of the *m* partial regression coefficients, and **R** is the conditional correlation matrix of the *m* expression levels. **R** was computed separately for *Hedonia* and *Eudaimonia* from the residuals of the multivariate linear model (equation 1) with all covariates in the model except the focal covariate. A *t* distribution with *n* – *m* degrees of freedom was used to test *t_Obrien_* against the null.

### Permutation *t*-test

The Fredrickson et al. 2013 [8] *t* test is similar to the O’Brien’s OLS test in that the test statistic is a composite of multiple *t*-values but uses an incorrect standard error. As an alternative to O’Briens OLS standard error, I used permutation to generate null distributions of the averaged *t*-values and then computed a *p*-values from this null distribution. To comply with the assumption of exchangeable error, I followed [2] and used the permutation method of [9]. For this procedure, the predictor variables were divided into main effects **Z** (hedonic and eudaimonic scores) and covariates **X** (the demographic and immune variables). Using the non-permuted data, the observed residuals (E_Y_) and predicted values (Ŷ) of **Y∣X** were estimated using equation 1. For each of the permuted iterations, rows of **E**_*Y*_ were permuted and added to the non-permuted Ŷ to generate a permuted response 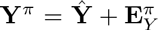, where *π* indicates a permuted value. Prior to fitting the permuted data, *Hedonia* and *Eudaimonia* and the *m* expression levels were re-centered and variance-standardized. The *m* coefficients and associated *t*-values for both *Hedonia* and *Eudaimonia* were then computed from the model **Y**^π^ **Y**∣**X** + **Z**. The test statistic is 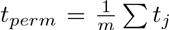, where *t_j_* is the *j*th *t*-value. 2000 iterations were run, including an iteration of nonpermuted data. The two-sided *p*-value of each hypothesis was computed as the fraction of ∣*t_perm_*∣ ≥ the observed ∣*t_perm_*∣.

### Rotation *z*-test (ROAST)

Because it is implemented in the function roast from the limma package [22], the rotation-test described in [25] is an attractive alternative to O’Brien’s test and the permutation *t*-test. The test statistic, *z_rot_*, is a mean *z*-score computed from the set of *m* moderated *t*-statistics computed for each gene. Using a hierarchical model, the moderated *t*-statistic uses information on the error of all genes in the set to estimate the gene specific standard error. A *p*-value for the test statistic is evaluated in a very similar manner to that described above in “Permutation test” but with some key differences. First, in the rotation-test, the observed residuals (**E***_Y_*) are from Y∣X where **X** includes not only the covariates but also the non-focal happiness score (for example, *Eudaimonia* is included in **X** for the test of *Hedonia*). Second, instead of permutation, the *n*-vector of residuals is rotated by a random vector *r*, which is constant for all genes within each iteration but variable among iterations. And third, the rotated residuals 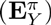 are used to directly compute the new *t*-statistics and *z*-scores without fitting the new model 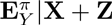 (where *π* indicates rotated residuals). The observed and rotated *z*-scores from 1999 rotations were used to generate the null distribution. The *p*-value for the “UpOrDown” test was used as this is the test of the two-tailed directional hypothesis. ROAST is small sample exact while the permutation of [9] is only asymptotically valid.

## Inference using linear model with fixed effects and correlated error

The model used by Fredrickson et al. 2015 [7] is

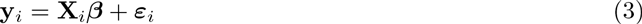

where **y***_i_* is the vector of *m* responses for subject *i*, ***X**_i_* is the model matrix for subject *i*, which includes the main effect *Gene* to identify the *j*th element of *y_j_*, and is the vector of coefficients, including the common effects of each covariate on the response. In this model, *ɛ_i_* ˜ *N(0, Σ)*, where *Σ* is the within subject error covariance matrix representing the correlated errors. The correlated errors result from random effects but the model does not explicitly model these. To implement this model, the data matrix with separate columns for each gene is stacked into long format by combining the *m* expression levels into a single response variable *(Expression)* and the variable *Gene* is created to identify the gene associated with a specific response (expression value). The univariate regression of *Expression* on the set of predictors results in the same OLS estimates as in the multivariate model described above. These estimates are unbiased but the standard errors for the estimates are incorrect because of the correlated errors. As in the multivariate model for the combined data, the model matrix includes a dummy variable indicating dataset (2013 or 2015).

### Generalized Estimating Equations

Fredrickson et al. 2015 [7] used GLS with a heterogenous compound symmetry error matrix to estimate the effects in Equation 3. Because only the fixed effects are of interest, I used Generalized Estimating Equations (GEE) with an exchangeable error matrix to estimate the fixed effects using the function geeglm in the geepack package [14]. The default sandwich estimator was used to compute the standard errors of the effects, which is robust to error covariance misspecification [16]. Nevertheless, GEE is less efficient if the error covariance is misspecified [23].

### Permutation and bootstrap GLS

Exploration of the behavior of the GLS as implemented by Fredrickson et al. 2015 [7] suggested partial regression coefficients that were more unstable than implied by the standard error. To explore the consequences of this instability on inference, I implemented both a bootstrap procedure to compute approximate standard errors and the Freedman and Lane permutation procedure [2] described above to compute permutation-GLS *p*-values. Each iteration of either the bootstrap or the permutation, the data were resampled (or the residuals permuted) in wide format, rescaled, and reshaped to long format. Coefficients were estimated using the gls function from the nmle package [20] using a heterogenous compound symmetry error matrix and the maximum likelihood method. The first iteration used the observed (not resampled) data. The standard partial regression coefficients and associated *t*-values for *Hedonia* and *Eudaimonia* were saved each iteration and used to generate standard errors for the bootstrap and a null distribution of *t*-values for the permutation. Because the time required to fit the GLS, and the exploratory goal of this analysis, I used only 299 iterations, which is sufficient for approximate, exploratory values. The regressor *Smoke* was excluded from the bootstrap analysis because some bootstrap samples had zero cases with level *Smoke* = 1, which leads to an unsolvable model. Because of this slightly different specification, I limited the bootstrap analysis to the FRED15 dataset. The GLS coefficients for FRED15 with and without *Smoke* in the model are 0.086 and 0.032 for *Hedonia* and −0.511 and −0.568 for *Eudaimonia*.

## Type I error, power, and exaggeration ratio

I used Monte Carlo simulation to explore type I, type II, and type M errors with data similar in structure to the Fred15 dataset. Type M error is error in the magnitude of the estimated coefficient when *p <*0.05 [10]. For type M error, I used the exaggeration ratio 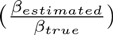 [10]. In each run of the simulation, a random *n* × *p* matrix X of independent variables (*n* samples of *p* covariates) and a random *n* × *m* matrix **Y** of response variables (*n* samples of *m* responses) were generated using the function rmvnorm from the mvtnorm package [11]. All simulated independent variables were modeled as continuous variables sampled from 𝒩**(0, S***_X_*), where **S***_X_* is the covariance matrix of the 17 regressor variable from FRED15. The 52 response variables were modeled as continuous variables sampled from 𝒩(**0, S***_Y_*), where **S***_Y_* is the covariance matrix of the 52 gene expression levels from FRED15. For the power simulations (including type M errors), the standardized effect of *Eudaimonia* on the mean response was set to 0.067, which is the estimated, standardized effect for the FRED15 dataset. The effect of all other covariates, including that of *Hedonia* was set to zero. For the type I simulations, all effects were set to zero. To explore the consequences of increasing *n* or increasing *m* on error rates, the simulation was run with three combinations of *n* and *m* (*n* = 122, *m* = 10; *n* = 22, *m* = 30; *n* = 366, *m* = 10). The *m × m* covariance matrix used to generate **Y** using the rmvnorm function was a random sample of **S***_Y_* each iteration.

**Table 1:**
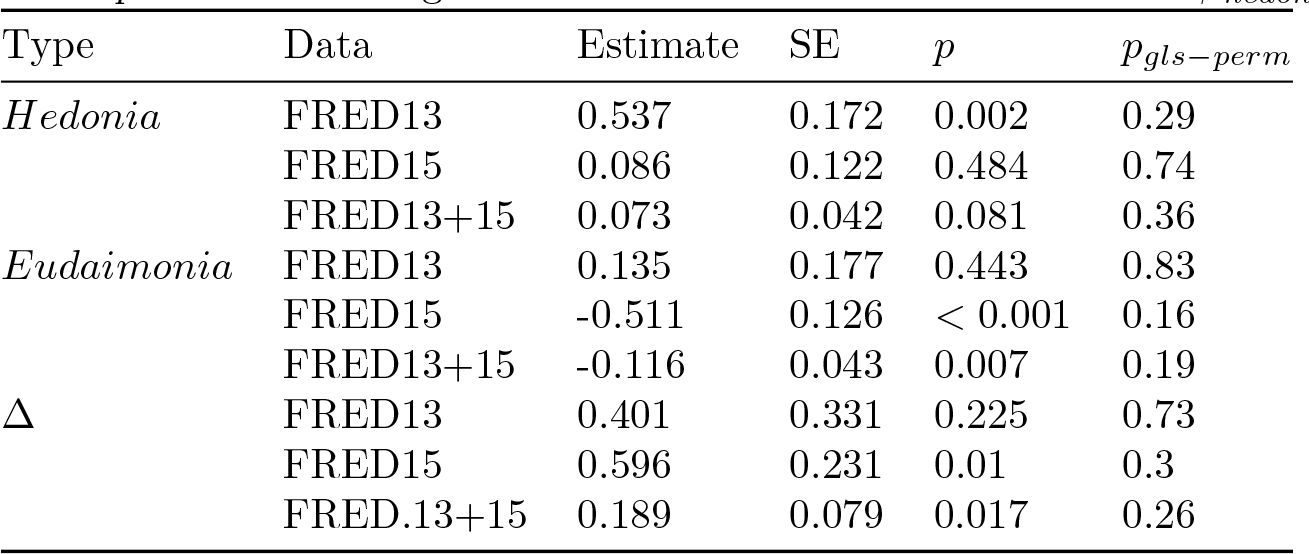
GLS estimates of the variance-standardized coefficients for the 2013, 2015, and combined data. The GLS-permutation *p*-values are also given. Δ is the difference in the estimates: *β_hedonia_ – β_eudaimonia_*

| Type | Data | Estimate | SE | $p$ | $p_{gls-perm}$ |
| --- | --- | --- | --- | --- | --- |
| *Hedonia* | FRED13 | 0.537 | 0.172 | 0.002 | 0.29 |
|  | FRED15 | 0.086 | 0.122 | 0.484 | 0.74 |
|  | FRED13+15 | 0.073 | 0.042 | 0.081 | 0.36 |
| *Eudaimonia* | FRED13 | 0.135 | 0.177 | 0.443 | 0.83 |
|  | FRED15 | -0.511 | 0.126 | < 0.001 | 0.16 |
|  | FRED13+15 | -0.116 | 0.043 | 0.007 | 0.19 |
| $\Delta$ | FRED13 | 0.401 | 0.331 | 0.225 | 0.73 |
|  | FRED15 | 0.596 | 0.231 | 0.01 | 0.3 |
|  | FRED.13+15 | 0.189 | 0.079 | 0.017 | 0.26 |

## Correlated estimation error

Regressors with a high positive correlation, as with *Hedonia* and *Eudaimonia*, have negatively correlated partial regression coefficients. I give a brief mathematical explanation of this in the discussion but also show this empirically using Monte Carlo simulation. The simulation was implemented precisely as described for the type I simulation above, except that I simulated all *m* = 52 gene expression levels in the FRED15 dataset. Each run of the simulation, the coefficients for *Hedonia* and *Eudaimonia* were estimating using GLS, OLS (multivariate), and GEE. 100 iterations were run to generate 100 pairs of points for the correlation.

All analyses were performed using R [21]. All data cleaning and analysis scripts are available at the public GitHub repository https://github.com/middleprofessor/happiness.

## Results

### Replication of previous analyses

The variance-standardized effects and *p*-values for hedonic and eudaimonic scores estimated from the GLS for each dataset are given in Table 1. My estimates for FRED15 and FRED13+15 are within 0.002 standard units of those reported in Fredrickson et al. 2015 [7]. Again, the coefficients for FRED13+15 reported here are from a model that did not include *IL*6 in the response as discussed in methods. Fredrickson et al. 2015 [7] do not report the GLS results for the 2013 data alone. My estimates of the coefficients for the FRED13 data show a pattern opposite to that for FRED15. That is, with the 2013 data, the effect of *Hedonia* is large and has a very small *p*-value (0.002) while the effect for *Eudaimonia* is small and not statistically significant (*p* = 0.44). My FRED13 coefficients are the same as those reported in the exploratory re-analysis of the FRED13 and FRED15 datasets by [4], who also note the opposite pattern from the 2015 results.

**Table 2:** OLS estimates of mean effects (Estimate) on CTRA gene expression. The estimates are the mean variance-standardized partial regression coefficients from the multivariate regression over the *m* responses (genes). Δ is the difference in mean effect. The SEs were estimated using a bootstrap. The *p*-values are from the bootstrap, O’Brien’s OLS, permutation *t*, and rotation *z* tests.

| Type | Data | Estimate | SE | $p_{boot}$ | $p_{obrien}$ | $p_{perm}$ | $p_{rot}$ |
| --- | --- | --- | --- | --- | --- | --- | --- |
| *Hedonia* | FRED13 | 0.026 | 0.118 | 0.83 | 0.8 | 0.81 | 0.78 |
|  | FRED15 | 0.062 | 0.044 | 0.16 | 0.24 | 0.23 | 0.23 |
|  | FRED13+15 | 0.052 | 0.041 | 0.21 | 0.22 | 0.2 | 0.22 |
| *Eudaimonia* | FRED13 | -0.063 | 0.125 | 0.62 | 0.5 | 0.49 | 0.49 |
|  | FRED15 | -0.067 | 0.048 | 0.17 | 0.2 | 0.18 | 0.2 |
|  | FRED13+15 | -0.064 | 0.038 | 0.1 | 0.14 | 0.13 | 0.13 |
| $\Delta$ | FRED13 | 0.089 | 0.235 | 0.71 | 0.64 | 0.63 | 0.61 |
|  | FRED15 | 0.129 | 0.085 | 0.13 | 0.22 | 0.17 | 0.18 |
|  | FRED13+15 | 0.116 | 0.074 | 0.12 | 0.17 | 0.14 | 0.14 |

### New results

Standardized mean effects 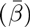 estimated by multivariate regression are very small and positive for *Hedonia* and very small and negative for *Eudaimonia* for both 2013 and 2015 datasets and the combined dataset (Table 2). The bootstrap SE for each mean indicates that a 95% confidence interval is too large to have any confidence in the direction of either of the effects for any dataset. The *p*-values computed using the bootstrap *t*-test, O’Brien’s OLS test, the permutation *t*-test, and the rotation *z*-test are very consistent with each other and all fail to reject the null. The OLS estimates of the difference (∆) between hedonic and eudaimonic effects are small and positive. The bootstrap SE for all Δ are too large to have any confidence in the direction of the difference and the *p*-values from each of the four tests fail to reject the null for any of the data sets. The GEE estimates are the same as the OLS estimates to the 2nd decimal place for all three datasets and the robust standard errors are similar to those estimated by bootstrap (Table 3). The GEE *p*-values are very similar to those from the OLS tests for all three datasets and fail to reject any of the nulls.

The GLS permutation *p*-values fail to reject the nulls for any of the tests (Table 1). Unlike the GEE *p*-values, the GLS permutation *p*-values are not similar to those from the OLS tests. The GLS bootstrap distributions of standardized effects for *Hedonia* and *Eudaimonia* for FRED15 are shown in Fig. 1. The standard errors of the effects computed from these distributions are 0.27 for *Hedonia* and 0.36 for *Eudaimonia,* which are 2–3 times the standard errors computed by the GLS model.

The GLS tests have inflated type I error rates that decreases with *n* but increases with *m* for a given *n* (Table 4). For the GEE test, type I error is moderately inflated (0.079 – 0.083) when the effective sample size is smaller (*n* = 122, *m* = 10 and *n* = 122, *m* = 30 but only slightly inflated when the effective sample size is larger (*n* = 366, *m* = 10). Type I errors for the Permutation *t*, the O’Brien OLS and the rotation *z*-score tests are close to the expected value of 0.05 for all three combinations of *n* and *m*. The GLS test has about 50% more power than the O’Brien or rotation tests when the effective sample size is low (*n* = 122, *m* = 1) but only about 5% more power than the O’Brien or rotation tests when the effective sample size is high (*n* = 366, *m* = 10). Of course this additional power for GLS comes at the expense of control of type I error. For all tests, power increases much more by adding subjects than by adding genes. The exaggeration ratio is about the same for all tests when effective sample size is low and high. Importantly, with increased *m* at a constant *n*, however, the exaggeration ratio is much higher for the GLS estimates than the OLS or GEE estimates.

**Table 3:** Generalized Estimating Equations estimates of the effects and difference in effects (Δ). SE is a robust standard error.

| Type | Data | Estimate | SE | $p$ |
| --- | --- | --- | --- | --- |
| *Hedonia* | FRED13 | 0.026 | 0.098 | 0.79 |
|  | FRED15 | 0.062 | 0.04 | 0.12 |
|  | FRED13+15 | 0.052 | 0.039 | 0.19 |
| *Eudaimonia* | FRED13 | -0.063 | 0.098 | 0.52 |
|  | FRED15 | -0.067 | 0.046 | 0.14 |
|  | FRED13+15 | -0.064 | 0.039 | 0.1 |
| $\Delta$ | FRED13 | 0.089 | 0.189 | 0.64 |
|  | FRED15 | 0.129 | 0.079 | 0.1 |
|  | FRED13+15 | 0.116 | 0.073 | 0.11 |

The expected, large negative correlation between the partial regression coefficients for *Hedonia* and *Eudaimonia* are shown using the GLS bootstrap distribution (Fig. 1) and using the GLS Monte Carlo simulation results (Fig. 2). Despite modeling the empirical correlations among the regressors and among the response variables, the distribution of standardized coefficients from the GLS Monte Carlo simulation have a much smaller range than that from the GLS permutation (e.g.95% confidence interval for *β_eudaimonia_* from the Monte Carlo simulation is −0.20 to 0.23 while that from the GLS permutation is −0.62 to 0.57), which suggests there is something about the structure of the actual data that is inflating the coefficient estimates [17]. Strongly biased estimates are also indicated by a diagnostic plot of residual versus fitted values from the GLS model (Fig. 3A). This bias is not apparent with the residual versus fitted values from the GEE model (Fig. 3B).

## Discussion

The re-analysis of the CTRA gene expression data in subjects scored for hedonic and eudaimonic happiness fails to support either the original conclusion of an opposite relationship of hedonic and eudaimonic happiness on the CTRA (conserved transcriptional response to social adversity) gene set [8] or the more recent emphasis on the large negative relationship between eudaimonic well-being and CTRA expression [7]. The consistency among the GEE test and the four OLS tests for each hypothesis and dataset is notable. A causal association between happiness components and CTRA expression levels would be an important discovery. Certainly, some association between happiness scores and CTRA expression levels must exist because of common shared paths within the complex network of causal relationships of the underlying physiology. Nevertheless, observational studies like that of [8] and [7] are poor designs for discovering knowledge [24].

**Table 4:** Type I error, power, and exaggeration ratio for the GLS, GEE, O’Brien’s OLS, Permutation *t*, and rotation *z* tests at different levels of number of subjects (*n*) and number of genes (*m*)

| model | $n$ | $m$ | iter | Type I | Power | ER |
| --- | --- | --- | --- | --- | --- | --- |
| GLS | 122 | 10 | 2000 | 0.109 | 0.29 | 2.4 |
| GEE | 122 | 10 | 8000 | 0.082 | 0.23 | 2.2 |
| O'Brien | 122 | 10 | 8000 | 0.05 | 0.19 | 2.3 |
| Perm $t$ | 122 | 10 | 3000 | 0.052 | 0.2 | 2.3 |
| ROAST | 122 | 10 | 8000 | 0.052 | 0.18 | 2.3 |
| GLS | 122 | 30 | 2000 | 0.152 | 0.37 | 2.4 |
| GEE | 122 | 30 | 8000 | 0.079 | 0.3 | 1.9 |
| O'Brien | 122 | 30 | 8000 | 0.049 | 0.25 | 2 |
| Perm $t$ | 122 | 30 | 3000 | 0.046 | 0.23 | 2 |
| ROAST | 122 | 30 | 8000 | 0.05 | 0.23 | 2 |
| GLS | 366 | 10 | 2000 | 0.084 | 0.54 | 1.5 |
| GEE | 366 | 10 | 8000 | 0.057 | 0.5 | 1.4 |
| O'Brien | 366 | 10 | 8000 | 0.048 | 0.49 | 1.4 |
| Perm $t$ | 366 | 10 | 3000 | 0.051 | 0.49 | 1.4 |
| ROAST | 366 | 10 | 8000 | 0.048 | 0.47 | 1.4 |

**Figure 1:**
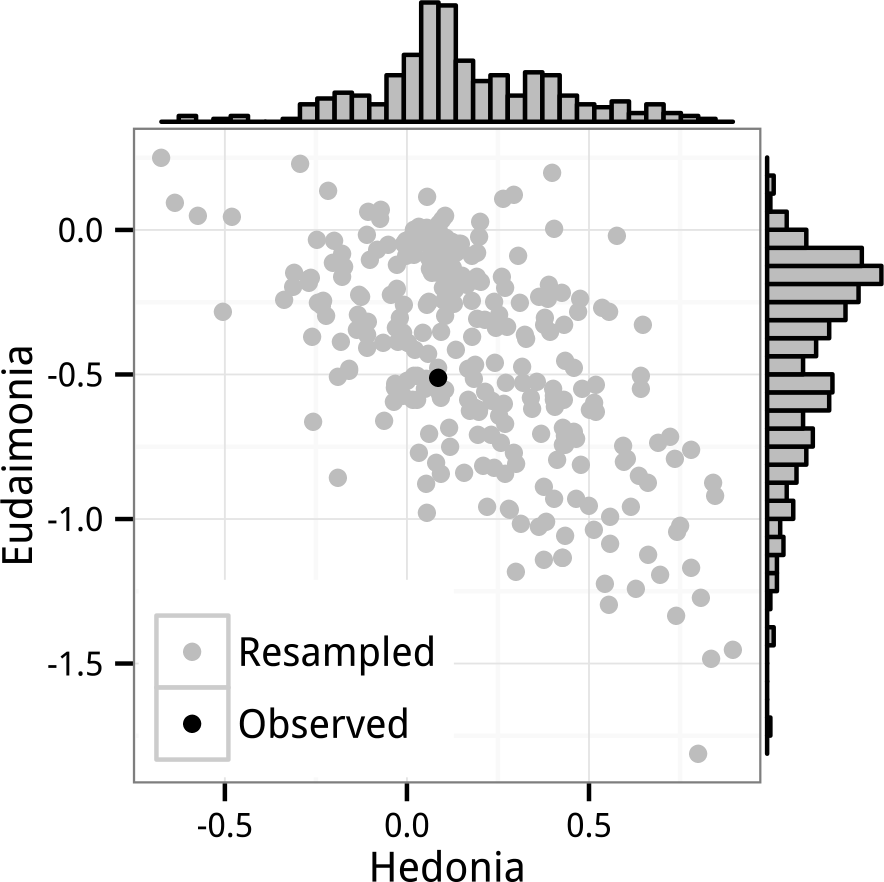
Distribution of GLS bootstrap resampled standard partial regression coefficients for *Hedonia* and *Eudaimonia.* The data are the FRED15 dataset and the coefficients were estimated by the linear model with correlated error (GLS). Also shown is the observed value for FRED15(black).

**Figure 2:**
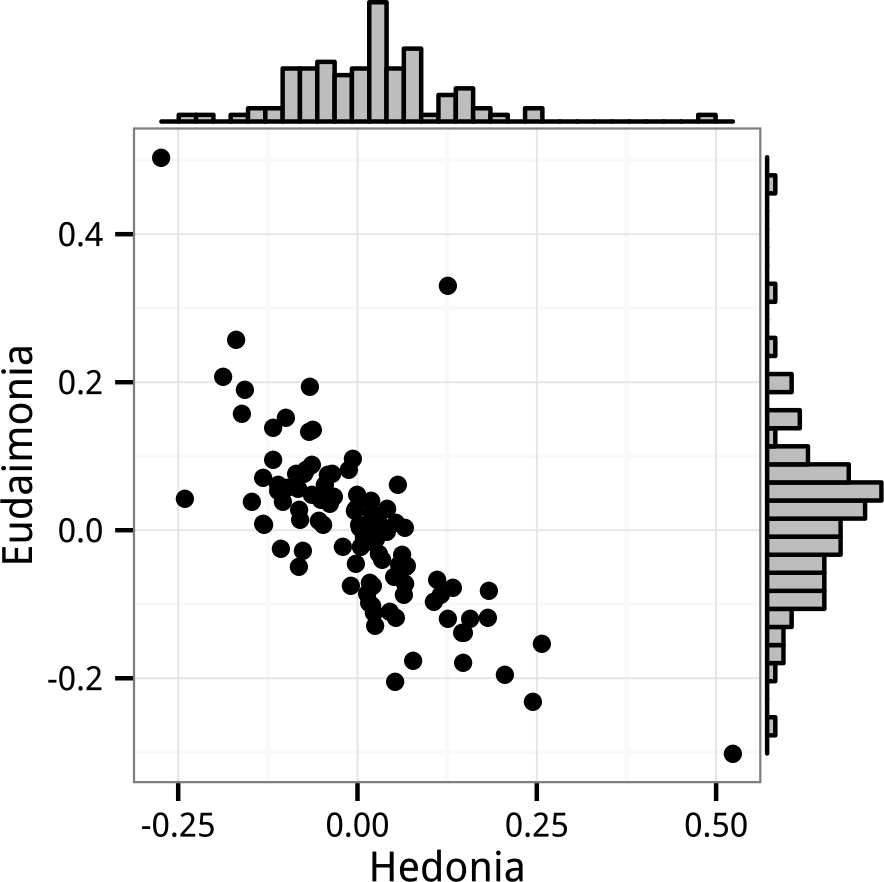
Bivariate distribution of standard partial regression coefficients for *Hedonia* and *Eudaimonia* from the Monte Carlo experiments. The Monte Carlo simulated the FRED15 data but with zero expected effect of any of the regressors on the gene expression levels. The coefficients were estimated by the linear model with correlated error (GLS).

The apparent replication of the sign of the effects between FRED13 and FRED15 [7] is consistent with the OLS estimates but strikingly inconsistent with the GLS estimates, although this failure of the GLS estimates to replicate was not noted by Fredrickson et al. 2015 [7] because they used the OLS estimates to illustrate the replication. Regardless, any replication in the sign of the mean effect should not be surprising given only two replicates of two coefficients. More importantly, as I show below, a pattern of coefficients of opposite sign is expected given the high positive correlation between hedonic and eudaimonic scores.

Several features of the original GLS results [7] suggest unstable and inflated coefficient estimates resulting from the GLS model. First, the opposite pattern of effects estimated in FRED13 and FRED15 suggests that either something very different biologically is going on between the subjects in FRED13 and FRED15 or the coefficients are more unstable than suggested by their (non-robust) standard error. Second, the GLS coefficients are very different from the OLS coefficients (Tables 1 and 2). Third, at least one of the GLS coefficients in each of the datasets is very large relative to what we’d expect from a gene set association given observational data and the stated hypotheses. Fourth, in a supplementary table, Fredrickson et al. 2015 [7] report strikingly different results (small, negative coefficients for both *Hedonia* and *Eudaimonia)* for the FRED15 dataset using an unstructured error matrix for the GLS computation *(Hedonia*: *β* = −0.014,*p* = 0.17; *Eudaimonia*: −0.0026, *p* = 0.81. Compare these to Table 1). Fredrickson et al. 2015 [7] failed to address any of these, including why the 2013 dataset was not analyzed using the updated (GLS) analysis or why the results using an unstructured error matrix were not the focus of the main paper.

The results reported here support the conclusion of inflated coefficient estimates. These results include the large coefficients that commonly occurred in the GLS with permuted data despite the expected effects of zero, the exaggeration ratio (computed from the Monte Carlo simulations) that increased with the number of outcome variables (*m*) relative to the number of subjects (*n*), and the diagnostic plot of the residual vs. fitted values that indicates biased estimates.

Downward biased standard errors in a GLS estimate with misspecified error matrix are well known, although the magnitude of the bias can be very specific to the degree of misspecification and effective sample size [15, 13]. The Monte-Carlo simulation results presented here, which modeled simulated data with the same covariance structure as both the predictors and responses in the FRED15 dataset, show that the GLS model results in increasingly inflated type I error when the number of responses (*m*) increases relative to the number of subjects (*n*). By contrast, the GEE, O’Brien’s OLS, permutation *t*, and rotation *z* tests maintain error rates close to the expected value (0.05) even with small *n* and even with increasing *m* relative to *n*.

The apparent replication of coefficients of opposite sign for *Hedonia* and *Eudaimonia* is consistent with trivially small effects (effectively equal to zero) in combination with the high empirical correlation between hedonic and eudaimonic scores (0.80 in FRED13 and 0.74 in FRED15). Partial regression coefficients of regressors that are positively correlated are themselves negatively correlated because their estimation shares common components that are of opposite sign. This is easily shown using the data from FRED15 where, disregarding all predictors but hedonic and eudaimonic scores, the partial regression coefficient of any gene expression level on *Hedonia* (*X*_1_) and *Eudaimonoia* (*X*_2_) are

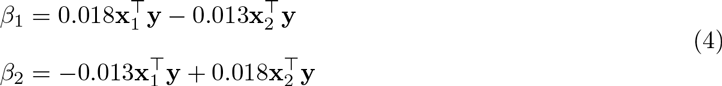

where the 0.018 and −0.013 are the diagonal and off-diagonal elements of the inverse of the **X**^T^**X** matrix of FRED15 (again disregarding all other predictors to simplify the explanation). Because of the high correlation between hedonic and eudaimonic scores, both *β_1_* and *β_2_*;include a large contribution from the covariance of the other *X* with *Y* but the sign of this contribution is negative. Consequently, if the true effects are trivially small, then the pair of *β* coefficients will tend to have opposite signs because of the negative correlation of estimates centered near zero. Random noise creates negatively correlated coefficients that tend to be opposite in sign. Linear mixed models do not adjust for this correlation. The negative correlation between coefficients is easily seen in the distribution of bootstrap GLS estimates of *β_hedonia_* and *β_eudaimonia_* (Fig. 1). The tendency for the coefficients to have opposite signs if the expected effects are zero is seen in the Monte Carlo simulation of the FRED15 data (Fig. 2). In conclusion, the most parsimonious explanation of the apparent replication of opposing effects of hedonic and eudaimonic scores on CTRA gene expression is correlated noise arising from the geometry of multiple regression.

**Figure 3:**
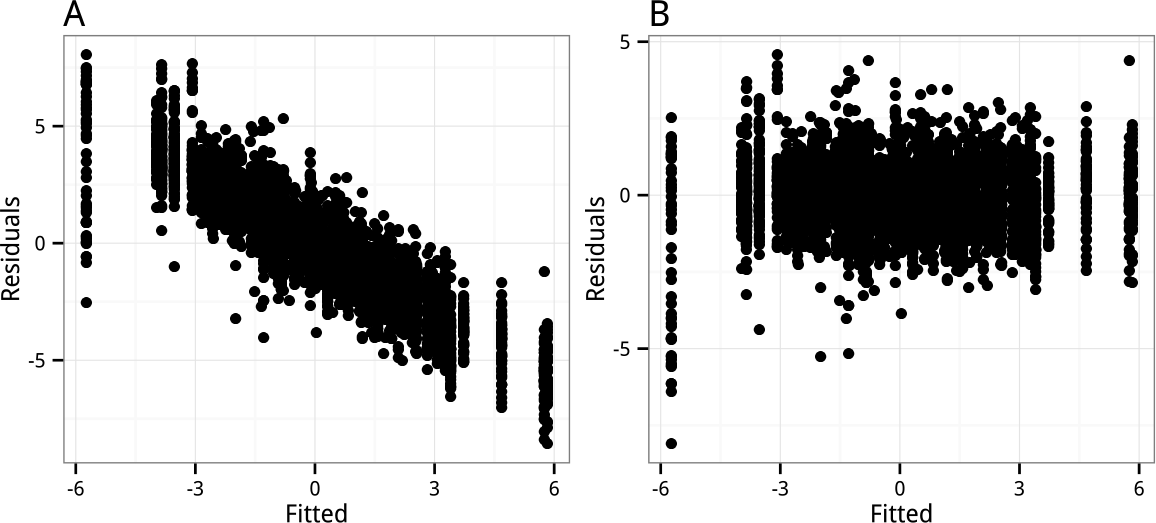
Residual versus fitted values from the A. fixed effects linear model with correlated error (GLS), and B. Generalized Estimating Equations (GEE) model. The pattern in *A* indicates strongly biased estimates.

